# Optimal thermal niche-tracking buffers wild great tits against climate change

**DOI:** 10.1101/2024.07.18.603942

**Authors:** David López-Idiáquez, Ella F. Cole, Charlotte E. Regan, Ben C. Sheldon

## Abstract

Animal and plant populations are responding to climate change by shifting the timing of key seasonal events. However, while measuring responses of natural systems on human calendars provides a standard scale of measurement it reveals little about the underlying biological causes and mechanisms of change. Here, using data from a six decade longitudinal, individual-based, study of great tits *Parus major*, we show that changing the timing of breeding has enabled birds to track specific temperatures at different stages in reproduction. We show, further, that multiple measures of reproductive success are maximised at intermediate, stage-specific, temperatures that correspond to the temperatures tracked while climate has changed in recent decades. Finally, we show that tracking temperatures has allowed great tits to match reproduction with a key food source for nestling development, the winter moth *Operophtera brumata*. Consequently, phenotypic plasticity in reproductive timing has enabled great tits to track temporal changes in the environmental conditions that maximise individual reproductive success. Shifting our perspective from analysing the phenological timing of life history events under climate change to analysing changes relative to environmental gradients has the potential to shed new light on the causes, mechanisms, and consequences of these shifts.

## Introduction

Rapid climate change is altering environments across the globe and to persist, populations must respond to these changes. Research on long-term populations has uncovered plastic and genetic responses to climate change in traits such as morphology [1], ornamentation [2], and phenology [3]. These phenotypic changes, such as the general reduction in body size across taxa [4], have allowed populations to cope, at least to some extent, with the effects of climate change [5]. Changes in reproductive timing are one of the best-described consequences of climate change [6], and the advance in egg-laying date of insectivorous birds to track the peak of food during breeding serves as a textbook example of phenological responses to global warming [7, 8, 9, 10].Besides phenotypic changes *in situ*, an alternative ecological response to climate change is tracking the optimal climatic niche through shifts in species distribution ranges. In general, as temperatures have risen, there is good evidence that species have moved their distributions poleward, to higher altitudes, or to greater depths, in search of cooler environments [11, 12, 13, 14].

These two types of response could be viewed as equivalent for some populations, with changes in phenology (e.g. earlier timing) enabling populations to track their optimal temperatures, allowing them to maintain a stable thermal niche without the need of a shift in their distribution ranges. This idea has been supported by two studies, one in plants [15] and another in birds [16], both of which focused on large spatial scales and across species to show that phenological changes can enable species to maintain stable climatic niches. Socolar et al. (2017) reported that advancing reproductive timing by between 5 and 12 days over the last century in Californian bird communities buffered the approximately 1° C increase in the area over the same period. Such population responses can, in principle, be produced by multiple mechanisms (e.g. evolution, plasticity or redistribution of individuals), which cannot be disentangled using cross-sectional data collected across long time intervals. Individually-based longitudinal studies enable more direct dissection of the mechanisms that underlie such responses [17], including the direct assesment of the fitness consequences of varying temperatures, but we are unaware of any such studies of the thermal niche-tracking hypothesis.

In this study, we explore the role of phenological change as a mechanism to track optimal temperatures during breeding by analysing an individually based dataset comprising more than 12 000 breeding attempts within a great tit *(Parus major)* population from 1965 to 2023. In this population, as in many others of this species [18], laying date has advanced over two weeks since 1965 in response to warming, making it an ideal study system to tackle our question [8, 3]. We analyse temporal trends in temperature from two different perspectives. First, based on calendar dates, we use a fixed temporal interval to quantify the change in temperatures experienced during reproduction over the course of the study period. Second, we measure long-term temperature changes in five distinct time intervals defined by individual reproductive timing; each of these represents a different stage of bird reproduction, with potentially different environmental sensitivities [19, 20, 21]. Identifying significant temporal trends in the fixed intervals, but not in the intervals relative to individual reproductive timing, would imply that the observed advance in laying date has allowed this great tit population to maintain stable temperatures during reproduction. We then expand these analyses linking temperatures in the different reproductive stages with two measures of reproductive success to analyse whether thermal tracking has allowed great tits to maintain optimal temperatures during reproduction. Finally, we use 45 years of data on winter-moth, *Operophtera brumata*, phenology from the same study site to analyse the link between relative temperatures and resource availability, as a potential mechanistic link between relative temperature and reproductive success.

## Results

### Long-term stability in thermal environment

Local mean temperature increased by 1.88° C across the fixed interval from February 15 to June 5 – corresponding to the reproductive period of great tits (*see methods*) – between 1965 and 2023 (0.032±0.005 °C *yr*^*−*1^, *F*_1,57_=31.21, *p <* 0.001, n=59, Fig. 1). In contrast, when we defined time intervals based on individual reproductive timing, we found that mean temperature has remained very stable across all of five periods between 1965 and 2023 (Fig. 1, Table 1). To explore potential biases introduced by comparing intervals of different lengths (the fixed interval is substantially longer than the relative intervals) we complemented our analysis of the fixed interval by exploring temporal trends in temperature for all periods of between 8 and 15 consecutive days (i.e. maximum and minimum duration of the relative intervals; *see methods*) between February 15 and June 5. The average increase of temperature from these models is 1.79° C [95% CI: 1.75, 1.83], closely aligning with the results obtained when using the full interval (Fig. 1). This shows that the different patterns found in the fixed and relative approaches are not a byproduct of the different interval lengths.

**Table 1:** Results from the linear mixed models analysing the temporal (i.e. annual) trends in temperature in the five intervals defined relative to individual reproductive timing. Egg-laying represents the period between laying the first and last eggs and incubation the period from the end of egg-laying to hatching. The periods hatching, nestling and fledging capture the time between hatching and 24 days post-hatch in 8-day intervals. The models included female individual identity and year (as a categorical variable) as random effects.

| Reproductive Period | Estimate | SE | F | P |
| --- | --- | --- | --- | --- |
| *Egg-laying (n=12315)* |  |  |  |  |
| Year | 0.001 | 0.009 | $F_{1,57.1}=0.021$ | 0.884 |
| *Incubation (n=11643)* |  |  |  |  |
| Year | -0.007 | 0.009 | $F_{1,57.1}=0.646$ | 0.424 |
| *Hatching period (n=11643)* |  |  |  |  |
| Year | -0.012 | 0.010 | $F_{1,57.1}=1.313$ | 0.256 |
| *Nestling period (n=11643)* |  |  |  |  |
| Year | -0.006 | 0.010 | $F_{1,57.1}=0.456$ | 0.502 |
| *Fledging period (n=11643)* |  |  |  |  |
| Year | -0.005 | 0.011 | $F_{1,57.0}=0.281$ | 0.597 |

**Figure 1:**
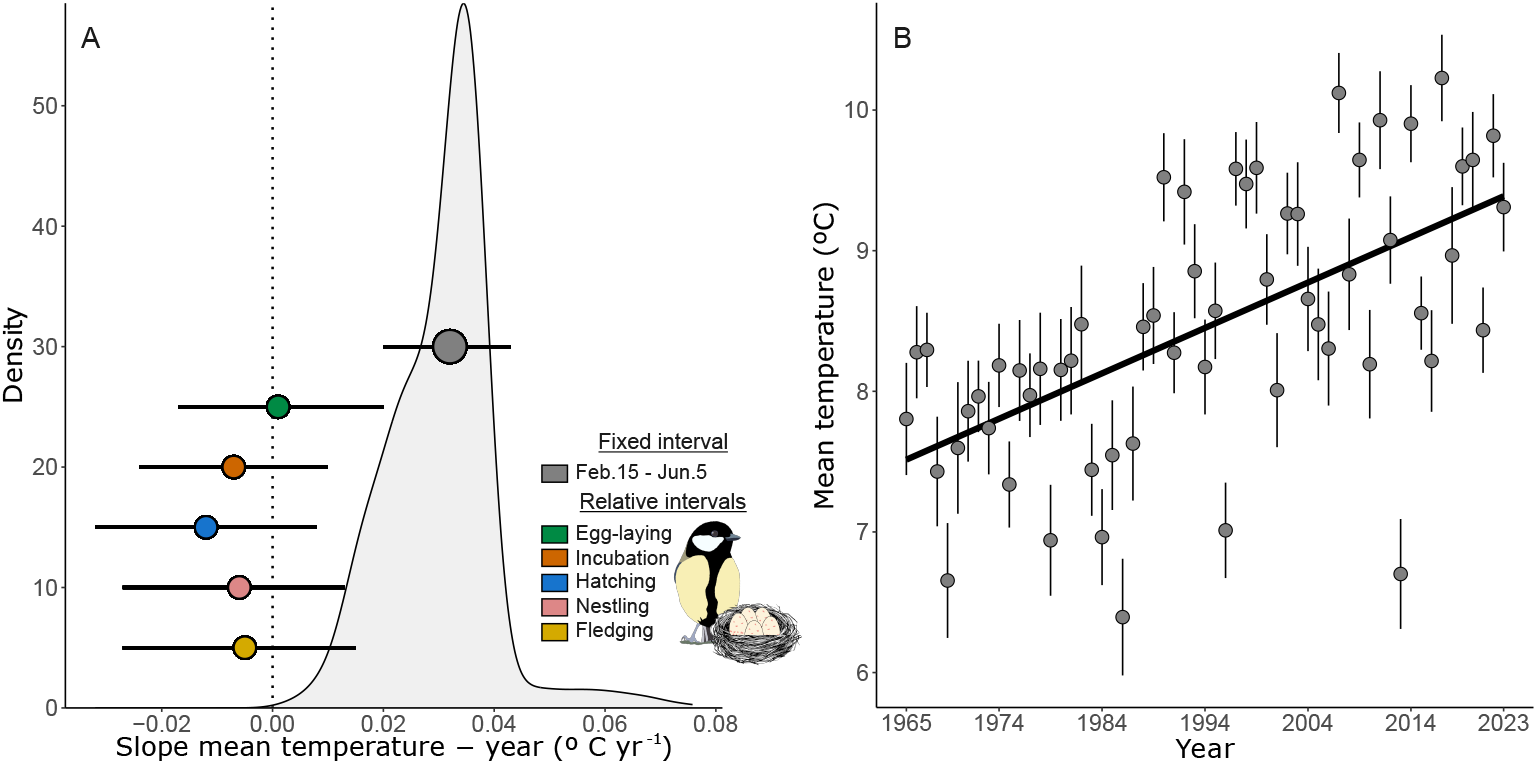
Temporal trends in temperature at Wytham woods near Oxford (UK). In A, the grey dot represents the slope of the temporal trend in mean temperature between 15 February and 5 June, and the coloured dots the slopes of the trends in mean temperature during empirically-defined egg-laying, incubation, and the hatching, nestling and fledging periods. The whiskers represent the ±95% confidence intervals (CI). 95% CIs crossing the line at zero denote non-significant associations. The density plot represents the distribution of the slopes computed for all intervals of between 8 and 15 consecutive days between the 15^th^ of February and 5^th^ of June. B represents the temporal trends in mean temperature (° C) in the interval between the 15^th^ of February and 5^th^ of June between 1965 and 2023. Dots and whiskers represent the yearly mean temperature ± standard error.

### Links between the thermal environment and reproductive success

We analysed the effects of mean temperature during the five relative reproductive stages on two measures of reproductive success: (i) the number of fledglings produced (analysed using hurdle models), and (ii) the proportion of eggs laid that were fledged. For fledgling number, the *conditional* component of the hurdle models, which models the non-zero part of the data, showed a strong negative quadratic association between number of fledglings and average temperature at egg-laying (−0.392±0.039, z=−9.88, *p <* 0.001). The number of fledglings decreased with lower and higher temperatures than 10.6° C during egg-laying (Fig. 2D). Number of fledglings also showed significant negative quadratic associations with mean temperatures at the hatching (−0.141±0.043, z=3.23, *p* = 0.001) and fledging (−0.175±0.058, z=−3.02, *p* = 0.002) periods, although these associations were less strong than with temperature in the egg-laying period (Figs. 2I, L). Mean temperatures in the remaining two periods (incubation and nestling) were not significantly associated with number of fledglings (*p >* 0.173, Fig. 2A; see supplementary material (SM)-1 for further detail on model results).

**Figure 2:**
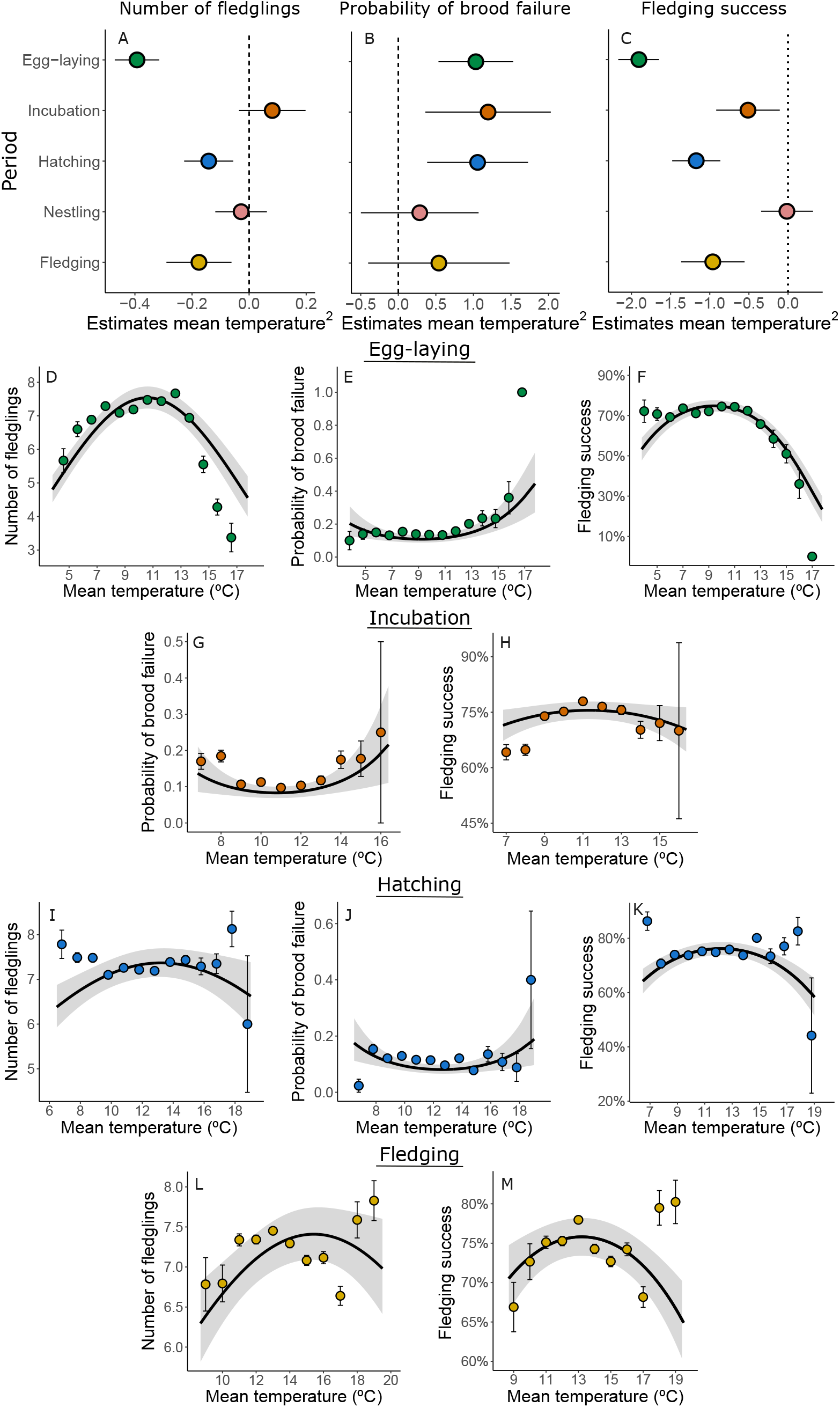
Associations between mean temperature (° C) during the five relative periods and reproductive success. A-C) Forest plots representing the estimates (±95% CIs) from the quadratic term of mean temperature for each studied period and reproductive measure. 95% CIs crossing the line at zero denote non-significant associations. The scatterplots represent the associations between mean temperature and reproductive success. The solid lines and grey ribbons represent the predicted association and 95% confidence interval from the model. The coloured dots and lines represent the average brood failure probability ± standard error grouped in 1° C bins.

The *zero-inflation* component of the hurdle model, which models the probability of an observation being zero (i.e. that a brood fails), showed quadratic positive associations with mean temperature during egglaying (1.032±0.253, z=4.067, *p <* 0.001), incubation (1.195±0.426, z=2.806, *p* = 0.005) and the hatching period (1.055±0.342, z=3.081, *p* = 0.002, Figs. 2E, J, The pattern was stronger during egg-laying and incubation than for the hatching period and in both cases the predicted trend suggests that the probability for a brood to fail increases with temperatures above 9.2 and 10.7° C respectively with weaker trends for lower temperatures. Temperatures during the nestling and fledging periods were not significantly associated with brood failure probability (*p >* 0.261, Fig. 2B, see SM-1 for further detail on model results).

Analysis of an alternative measure of reproductive success, fledging success (the ratio of eggs that become fledglings, which measures the efficiency of reproduction), also revealed significant negative quadratic associations with temperature during specific reproductive periods: egg-laying (−1.905±0.131, z=−14.505, *p <* 0.001) and the incubation (−0.511±0.207, z=−2.467, p=0.013), hatching (−1.172±0.156, z=−7.463, *p <* 0.001) and fledging periods (−0.961±0.207, z=−4.643, *p <* 0.001). The predicted models show that the pattern is stronger for temperature at egg-laying where fledging success decreased at lower, but particularly at higher temperatures than 9.4° C (Fig. 2F). In the three other periods with significant associations, the patterns were less strong than for egg-laying, but still showed that breeding at higher and lower temperatures than optimum was associated with reductions in fledging success (Fig. 2H, K, M). Temperature during the nestling period was not significantly associated with fledging success (p=0.931, Fig. 2C, see SM-2 for further detail on model results).

### Population tracking of the optimum temperatures

The frequent observation that reproductive success was maximised at intermediate stage-specific temperatures suggests the existence of an optimum temperature for reproduction. Therefore, we compared the optimum temperatures for maximising reproductive success in the relative periods with the median temperature in each period with the aim of assessing the extent to which our population has tracked optimum breeding temperatures (Fig. 3). A perfect match between the two would indicate that the phenological shift exhibited by the great tits in our population has allowed a close tracking of their optimal temperatures for reproduction. Although the sample size precludes formal analysis, Figure 3 supports this idea, particularly for the periods where temperature significantly influenced reproductive success.

**Figure 3:**
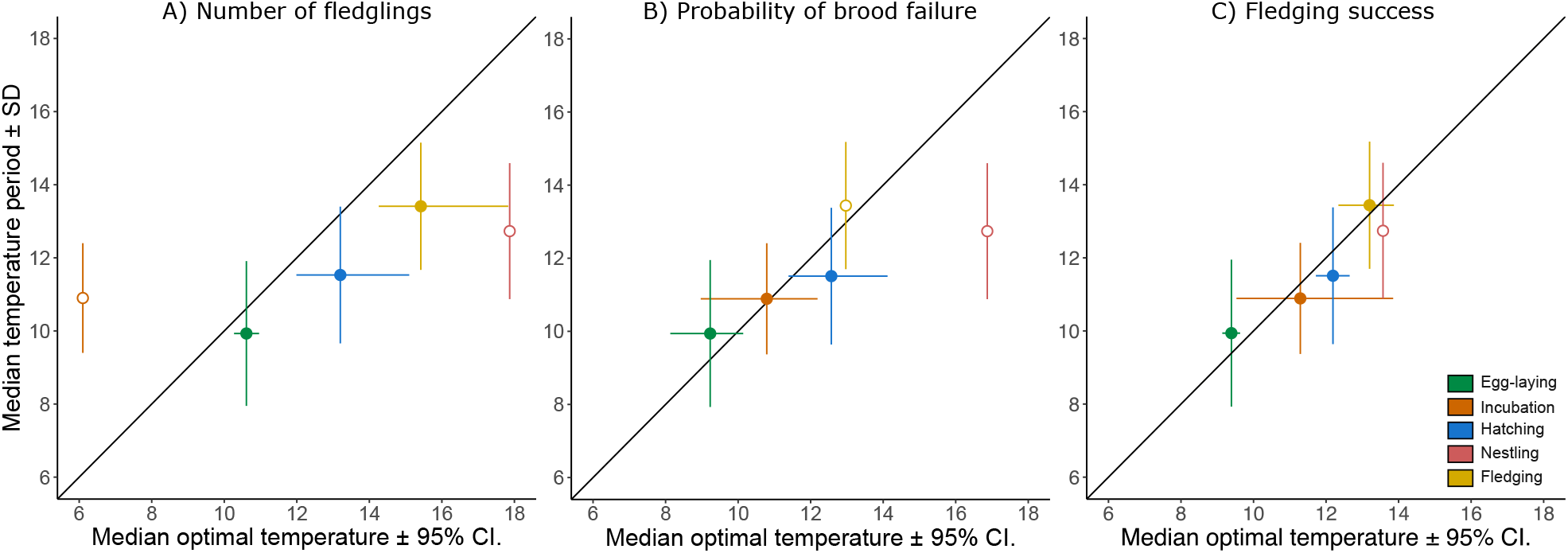
Representation of the median ±95% credible interval optimal temperature (°C) to maximise number of fledglings (A), minimise brood failure (B) and maximise fledging success (C) in each stage of reproduction, and the median average temperature observed in each period. Median optimal temperatures (and 95% credible intervals) were estimated by from the posterior distributions of Bayesian models fitted as those presented in the paper (see SM-3 for further information). For associations between reproductive success and temperature that were non-significant (empty dots) we do not represent the credible interval for the optimal temperature. The diagonal solid line (y = x) represents a perfect fit between median temperature in a period and temperature that maximises reproductive success.

### Thermal niche-tracking by winter moths

Analysis of standardised timing measures (half-fall date) from 45 years that were available between 1965-2023 showed that the seasonal phenology of winter moths has advanced by ∼16 days since 1965 (−0.270 ±0.056 days *yr*^*−*1^, *F*_1,43_=23.26, *p <* 0.001, n=45, Fig. 4A). As was the case for great tit phenology, temperature in the period around winter moth half-fall (±7 days) has remained stable between 1965 and 2023 (−0.002±0.011 °C *yr*^*−*1^, *F*_1,43_=0.071, *p* = 0.790, n=45, Fig. 4B).

**Figure 4:**
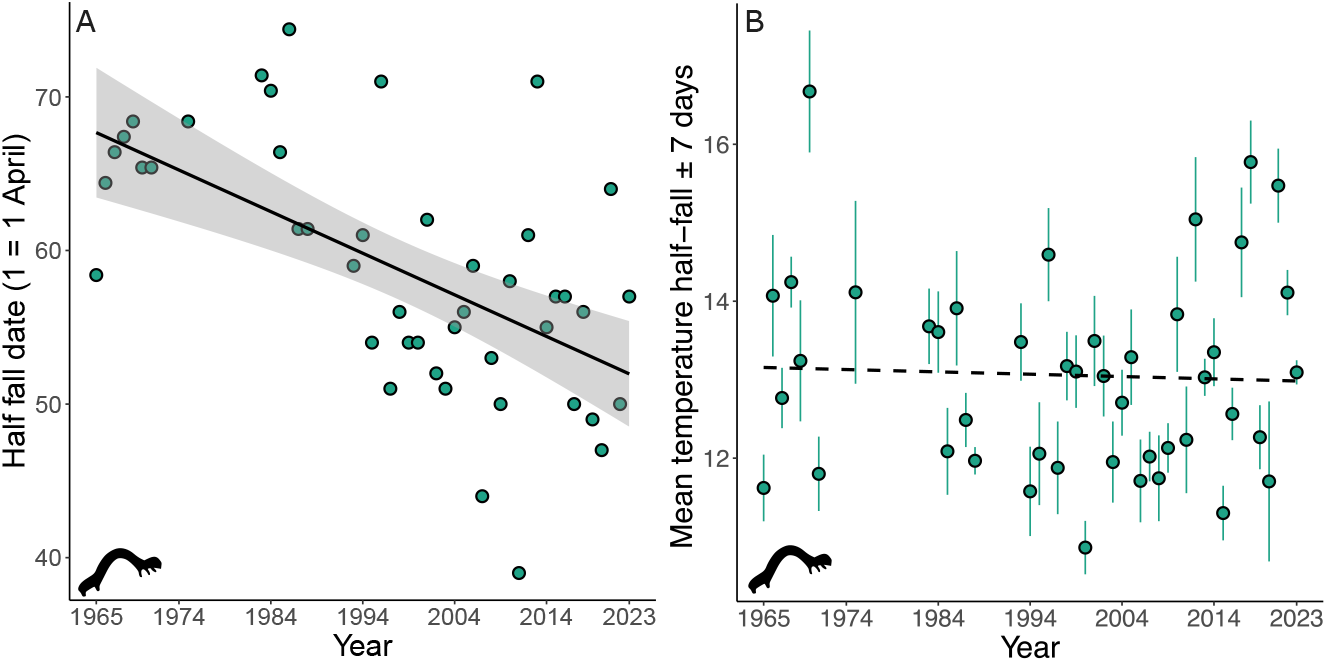
Temporal trends in half-fall (A) and in average temperature (° C) ± standard errors at half-fall ± 7days (B) by winter moths at Wytham woods. Solid lines represent significant and dashed non-significant trends.

### Links between thermal niche and resource overlap

Using (i) the individual-level data on breeding dates of great tits between 1965-2023, and (ii) annual data on standard measures of winter-moth timing, we calculated a measure of matching between timing of tit breeding and resource availability as: *match* = *hatching date + 10* – *half-fall date*. The rationale for this measure is that energetic demand of nestlings is maximized at around 10 days of age [22] and that late instar larvae provide the highest nutritional value. We then asked how individual-level temperature during specific reproductive periods related to the predicted match with the resource peak. We found positive quadratic patterns between temperature in all four reproductive periods (the fledging period was not included in this analysis as it frequently occurs after the caterpillar half-fall date) and absolute mismatch with winter moth caterpillar peak. During egg-laying (0.217±0.008, *F*_1,9486.1_=657.6, *p <* 0.001, Fig. 5A) and the hatching (0.165±0.009, *F*_1,9437.0_=302.1, *p <* 0.001, Fig. 5C) and nestling periods (0.266±0.009, *F*_1,9643.4_=845.08, *p <* 0.001, Fig. 5D) breeding at warmer and colder temperatures than optimal increased the mismatch with the peak of caterpillar availability. During incubation (0.256±0.015, *F*_1,8194.1_=258.3, *p <* 0.001, Fig. 5B), the quadratic pattern was mostly driven by the pre-peak trend meaning that incubating the clutch at lower rather than higher temperatures than optimum increases the mismatch with the peak of caterpillar availability (see SM-4 for further information on model results). Hence, these analyses are consistent with the suggestion that matching timing with a key trophic level is an important factor in driving the tight linkage between temperature and specific reproductive stages in great tits.

**Figure 5:**
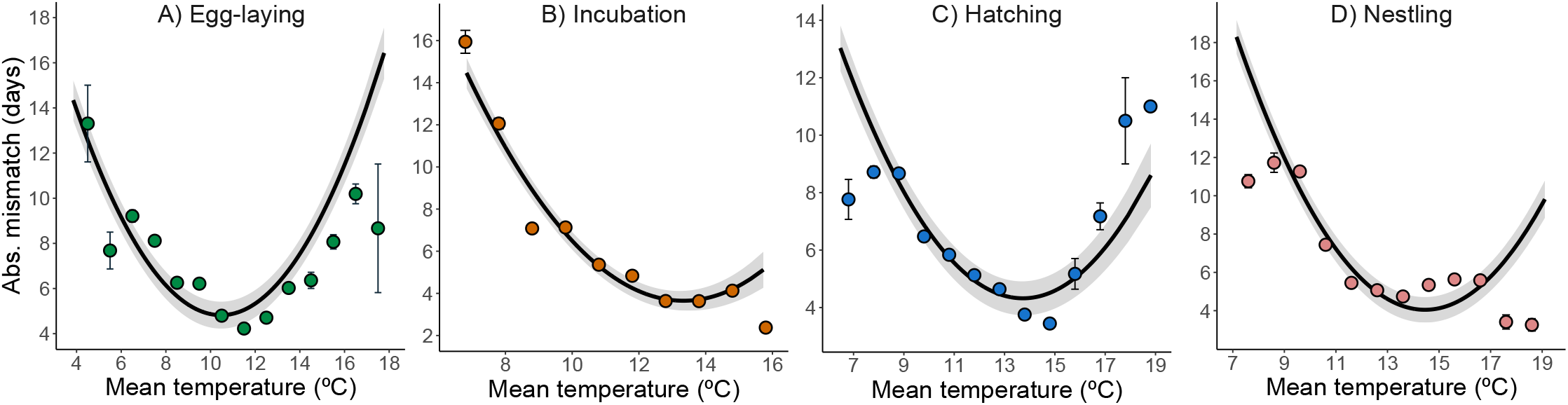
Associations between the absolute mismatch with the peak of food availability (half-fall date) and mean temperature (° C) in the intervals relative to individual breeding. The solid line and grey ribbon represent the predicted association and 95% confidence interval from the model. The coloured dots and lines represent the average mismatch ± standard error grouped in 1° C bins. The fledging period was not included as it usually occurs after the half-fall date.

## Discussion

Using an individually based dataset comprising over 12 000 breeding attempts, we demonstrate that the 15-day phenological advancement in timing of breeding over six decades in our population has allowed great tits to maintain stable temperatures during reproduction, buffering the almost 2° C increase in average temperature observed between 1965 and 2023. This finding supports the untested hypothesis that phenological adjustments enable populations to track their optimal thermal niches without the need of geographical range shifts. By tracking their optimal thermal niches great tits have maximised their reproductive success as thermal tracking has allowed them to synchronize with the peak of food availability. Overall, our results demonstrate that the stasis in temperature during breeding is consistent with natural selection acting to maintain a position at an optimum.

Phenological changes are a well-described response to climate change, with much of the existing research focusing on changes in reproductive timing measured relative to human calendar dates [23, 24]. However, the environmental consequences of changes in phenology have received less attention [15, 16]. Here, we show that the phenological advancement of our population has resulted in what is effectively thermal homeostasis during great tit reproduction, effectively conserving optimal thermal conditions despite rising temperatures (Fig. 3). This result confirms the pattern found by Socolar et al. (2017) for Californian avifauna comparing phenology and breeding temperatures in two time periods separated by over a century, but using individual-based phenotypic and fitness data at considerably higher spatial and temporal resolution. Our results also align with recent studies focusing on optimal reproductive timing [25, 10]. de Villemereuil et al. (2020) found that several mammal and bird populations exhibit fluctuations in the optimal reproductive timing, with relatively early or late breeding being favoured in different years. Our analyses suggest that these fluctuations probably represent an individual adjustment in reproductive timing as a response to the environmental change to breed at optimal temperatures, or in other words, that the changes in optimal reproductive timing are a consequence of tracking an optimal thermal niche during breeding. Youngflesh et al. (2023), analysing ringing data from 41 bird species across North America, found that breeding productivity decreased when breeding deviated from a phenological optimum. Our analyses linking temperatures during breeding to reproductive success add an individual-level analysis and mechanistic aspect to this general pattern. We found that most measures of reproductive success are maximised at intermediate breeding temperatures. Our results are, thus, consistent with natural selection having acted to enable tracking optimal temperatures during breeding, something that is highlighted by the close alignment of average temperatures in the periods relative to breeding with those that maximise reproductive success (Fig. 3).

Associations between temperature in the relative reproductive periods and reproductive success might, in principle, be explained by the direct effects of temperature on adults and nestlings. For instance, temperature experienced by the embryo during incubation and by nestlings and fledglings can directly impact factors such as metabolic and growth rates and also fledging success [26, 27] and survival [28, 29]. However, given that the bird population we studied is located in a temperate area with generally mild temperatures, and that great tits have wide geographic and thermal ranges, we expect that temperatures during reproduction, at least within normal ranges, do not impose a strong constraint. Rather, we consider it more likely that temperature has indirect effects on reproductive success by altering synchronisation with resource availability [30]. While tits can maintain some degree of synchronization through changes in hatching date [31, 32], the different effects of thermal environments on development rates of endotherms and exotherms are expected to lead to mismatch if temperatures are unusually cool or warm [7]. This could lead to greater mismatch with the peak of food demand by nestlings, thus, increasing the relative costs of reproduction for adults [33]. Our results validate the indirect effects of temperature on reproductive success, as we identified optimum temperatures in relative breeding periods that minimise the mismatch between the peak of food availability and demand. Overall, we show that breeding at below optimum temperatures during egg-laying, incubation and the hatching and nestling periods increased the mismatch with the caterpillar peak. Our results for egg-laying and for the hatching period also show that experiencing above optimum temperatures during those periods increased the mismatch. Given that low and high temperatures are expected to respectively delay, and advance, caterpillar emergence is possible that the increased mismatch occurs because of a limited capability of great tits to adjust the timing and development of breeding after the clutch has been laid.

The proximate mechanisms by which thermal tracking is achieved are not known. Research on reproductive timing of birds has explored two main cues used to time the start of reproduction, temperature and tree phenology. Experimental work has shown that captive great tits directly respond to changes in average temperature and warming rates [34, 35]. For instance, Visser et al. (2009) experimentally exposed great tits to temperature profiles from years with particularly early and late laying dates, with temperatures in the early treatment being higher than in the late treatment. They found that birds in the early treatment, which experienced warmer temperatures, laid earlier clutches than those in the late treatment. Laying dates from birds under the early treatment, however, differed from those that experienced the same temperature profiles in the wild, which suggests the need of extra cues in addition to temperature to time reproduction. Tree phenology, in this case tree green-up dates, is a likely additional cue that might be used by great tits to time the start of breeding ([36, 37, 38] but see [39, 40]). In our population, both temperature [8] and tree phenology [38, 41] have been identified as relevant factors modulating great tit reproductive timing. Thus, it is possible that the direct effects of temperature and indirect effects of phenology at lower trophic levels allow great tits to track optimal temperatures during breeding. Disentangling the role of these factors and others (i.e. insect phenology) in free-ranging populations will be a challenging task.

Further work could test whether the thermal niche tracking pattern found here can be extended to other species and contexts, as it provides an assessment of the extent to which populations are responding to climate change assessed against a more biologically relevant measure than calendar date. In our study, long-term data on caterpillar half-fall dates showed that the timing of this event has advanced by approximately 16 days between 1965 and 2023. Winter moth caterpillars face strong selection to synchronise their emergence to the availability of newly emerged leaves, as mismatch with tree budburst imposes strong fitness costs both in terms of survival and weight at pupation [42]. In this context, it would be expected that thanks to their phenological advancement, winter moths have been able to maintain stable temperatures as caterpillars, although we lack detailed long-term information about life stages other than the fifth instar reported here. Hence, these data suggest that thermal tracking may also occur in taxa other than birds, something that can be expected for the tri-trophic (birds, insects and trees) system we study.

Overall, our analyses show that the behavioural responses to climate warming in our study population have allowed great tits to maintain stable temperature during breeding. Viewing the birds’ decisions about when to breed as the choice of a temperature, rather than the choice of a date, offers the opportunity for a new perspective on the causes, mechanisms, and consequences of phenological responses to climate change. Scaling this perspective up to different populations and different species will reveal the generality of pattern described here. Scaling it down by using environmental data collected at smaller scales (e.g. at the territory level) has the potential to reveal novel understanding of the mechanisms behind phenotypic plasticity to changing environments. This perspective, of measuring responses against biologically relevant gradients, could be extended to other forms of environmental change.

## Methods

### Study system

We carried out this study in a nest box population of great tits at Wytham Woods (51°46N, 1°20W) a mixed deciduous woodland in Oxfordshire, which has been continuously monitored since 1947 [43]. Population monitoring includes nest box visits at least once a week during the breeding season (April-June), enabling the collection of detailed breeding data including laying date (date in which the first egg is laid), clutch size, hatching date, and fledgling success for each pair. Data on winter moth half-fall dates at Wytham were supplied by Dr L. Cole. Half-fall date is the day on which 50% of the seasonal total of winter moth caterpillars are collected in water traps under oak trees and provides a robust index of the peak of food availability for great tits [19, 31, 3]. In this study, as in previous work [21], we decided to define the starting year in 1965 to avoid potential biases introduced by the provision of new boxes up to 1961; mean generation time of a great tit is *<* 2 years [44]. Winter moth half-fall data was available for 45 of those 59 years (non-available years: 1972-1974, 1976-1982 and 1989-1992; analysis of great tit data remain qualitatively the same when using this reduced dataset, see SM-5 for further information).

### Temperature data and period definition

We obtained daily mean temperature data from the Central England dataset at the Met Office Hadley Centre https://www.metoffice.gov.uk/hadobs/. To analyse temporal trends in temperature between 1965 and 2023, we followed two approaches. First, we used a fixed interval spanning February 15 to June 5. February 15 is the beginning of the interval best explaining the onset of egg laying in our population, which was obtained using a sliding window analysis (see SM-6 for further information). To capture temperature across the whole breeding season we expanded the interval until the 5^th^ of June, which represents the average end of breeding in our population, computed as the addition of average laying date, average incubation duration, and 20 days (i.e. approximate time between hatching and fledgling). Second, we examined temperature changes in five periods relative to the observed timing of each individual reproductive event [19, 20, 21]. This included *egg-laying* (period from the first to the last egg), *incubation* (from the start of incubation to hatching), and the *hatching* (from hatching to seven days post-hatch), *nestling* (from eight to 15 days post-hatch), and *fledging* (from 16 to 21 days post-hatch) periods. We did this by matching daily meteorological data with each individual breeding attempt (see SM-7 for an illustrative figure). For instance, temperature prevailing at each individual breeding attempt during the egg-laying period was computed as the average temperature in the days between laying of the first and last eggs. These periods represent different stages in great tit breeding and likely show different sensitivities and responses to temperature changes in them. For instance, while during laying and incubation tits can adjust their reproductive timing to maximise the match with the peak of food availability, this cannot happen during the subsequent reproductive stages. In those, temperature may have direct and indirect effects on the brood, as the hatching period captures the stage when chicks are only starting to thermoregulate and the nestling period the stage when food requirements are highest.

### Statistical analyses

#### Temporal trends in temperature using fixed and relative intervals

To analyse the temporal trends in mean temperature on the fixed interval (from the 15^th^ of February to the 5^th^ of June) we fitted a linear model that included the yearly average temperature in that period as dependent variable and year (as a continuous variable) as fixed effect. To investigate the potential bias introduced by the selected length of this interval (e.g. longer intervals may be less prone to stochastic variation), we repeated this analysis with the same model structure for all intervals of between 8 and 15 consecutive days, which represent the minimum and maximum length of the relative intervals and thus, a more biologically relevant and comparable time frame than the whole interval.

To analyse the temporal trends in temperature in the relative periods we fitted one linear mixed model (LMM) per period. In these models, mean temperature in each individual breeding attempt and period was included as dependent variable and year (as a continuous variable) as a fixed effect in all models. To control for the nonindependence of data taken on the same years and from the same individuals we included year (as categorical variable) and female identity as random effects.

#### Association between temperature during the relative intervals and reproductive success

We analysed the association between temperature in the relative periods and two proxies of breeding success: number of fledglings and fledgling success (i.e. ratio of eggs that become fledglings).

For number of fledglings, we fitted hurdle models, which are employed to model data that is zero-inflated. These models have two parts, the conditional part that models the non-zero counts and that in our case assumed a Poisson error distribution and the zero-inflated part that models the probability of an observation being zero (i.e. the probability of having a failed brood). We ran one model per period that included number of fledglings as dependent variable and mean temperature of the period and its quadratic component as fixed effects in addition laying date to control for the well-known effects of reproductive timing on clutch size [43], and number of neighbours to control for the potential effects of competition/density on reproductive success, as covariates.

For fledgling success, we fitted generalized linear mixed models (GLMM) with a binomial distribution of errors. These models included fledgling success (i.e. proportion of eggs that become fledglings) as dependent variable and mean temperature of the period and its quadratic component as fixed effects in addition to laying date, clutch size, and number of neighbours.

In these models, year (as a categorical variable) and female identity were included as random effects, and all fixed effects were scaled to a mean of zero and standard deviation of one.

#### Temporal trends in half-fall dates and thermal niche tracking in winter moths

We analysed the temporal trends in half-fall dates by fitting a linear model with half-fall date (as number of days since the first of April) as dependent variable and year (as a continuous variable) as fixed effect. To analyse the temporal trends in temperature at half-fall date we computed the yearly average temperature ±7 days around half-fall and included this information as dependent variable in a linear model with year (as a continuous variable) as fixed effect. To explore how the selection of the intervals length influenced our results we repeated this analysis using ±15-, and 20-day intervals, with consistent results in all of them (see SM-8 for further information).

#### Temperature during the relative periods and mismatch with peak of caterpillar availability

We computed the mismatch between the peak of food demand and the peak of food availability as the absolute number of days between the date when nestlings are 10 days old (i.e. peak of food demand) and half-fall date [7]. To analyse the link between temperatures at the relative periods and mismatch we fitted one linear mixed model per period that included absolute mismatch as dependent variable and the temperature of the period and its quadratic component as explanatory. In these models we included year (as a categorical variable) and female identity as random effects. In this set of analyses, we did not include the fledging period, as it occurs mostly after the peak of food availability. Note that the sample size differences in this set of models is caused by missing half-fall data in some years. Using the raw instead of the absolute mismatch values yields qualitatively the same results (see SM-9 for further information).

All models were fitted using R (v.4.3.1; [45]) and the packages *lmerTest* (v.3.1-3; [46]) to fit the linear models, LMMs and GLMMs and *glmmTMB* (v.1.1.7; [47]) to fit the hurdle models. In the linear models we visually inspected the distribution of model residuals, not finding strong evidence for deviations from normality. In the models with *poisson* and *binomial* distributions we computed overdispersion values which in all cases were low *<* 1.39. The predicted trends shown in the plots were obtained using the package *ggeffects* (v.1.3.1; [48]) and the 95% C.I. were obtained using the package *confintr* (v.1.0.2; [49]).

## Acknowledgements

We thank all researchers and fieldworkers involved in the long-term monitoring of Wytham’s great tit population. The work was funded by NERC grant NE/X000184/1 and UKRI Frontiers grant EP/X024520/1, and recent data collection for the long-term study by BBSRC (BB/L006081/1), ERC (AdG250164) and NERC (NE/K006274/1 and NE/S010335/1).We are grateful to N. Merino Recalde and J.P. Woodman for analytical advice, and to L. Cole for providing data on winter moth half-fall dates.

## Author contributions

DL-I and BCS conceptualised the idea. DL-I conducted the data analysis with input from all authors. DL-I produced the first draft of the manuscript and all authors provided comments and contributed critically to the development of the final version.

