## Supplementary material (SM) for "Optimal thermal niche-tracking buffers wild great tits against climate change"

**Supplementary Material 1**

*Outputs from the models linking average temperature in the periods relative to breeding and number of fledglings.*

SM-1 Table 1: Results from the hurdle models analysing the association between number of fledglings and temperature at egg-laying. The *conditional part* models the non-zero counts of our data and the *zero-inflated part* that models the probability of an observation being zero (i.e. the probability of brood failure). Bold denotes statistical significance. The model included mother identity and year as random factors.

| Variable | Estimate | Std. Error | z | P |
|---|---|---|---|---|
| *Temperatures at egg-laying (n=11033)* | | | | |
| Conditional model | | | | |
| **Mean Temp.** | **0.416** | **0.039** | **10.52** | **<0.001** |
| **Mean Temp^2.** | **-0.392** | **0.039** | **-9.88** | **<0.001** |
| **Laying date** | **-0.099** | **0.007** | **-13.29** | **<0.001** |
| **Num. Neighbours** | **0.013** | **0.003** | **3.38** | **<0.001** |
| *Zero inflation model* | | | | |
| **Min Temp.** | **-0.950** | **0.262** | **-3.623** | **<0.001** |
| **Min Temp^2.** | **1.032** | **0.253** | **4.067** | **<0.001** |
| Laying date | 0.055 | 0.051 | 1.086 | 0.277 |
| Num. Neighbours | -0.010 | 0.029 | -0.354 | 0.723 |

SM-1 Table 2: Results from the hurdle models analysing the association between number of fledglings and temperature at incubation. The *conditional part* models the non-zero counts of our data and the *zero-inflated part* that models the probability of an observation being zero (i.e. the probability of brood failure). Bold denotes statistical significance. The model included mother identity and year as random factors.

| Variable | Estimate | Std. Error | z | P |
|---|---|---|---|---|
| *Temperatures at incubation (n=10421)* | | | | |
| Conditional model | | | | |
| Mean Temp. | -0.039 | 0.060 | -0.65 | 0.516 |
| Mean Temp.^2 | 0.081 | 0.059 | 1.36 | 0.173 |
| **Laying date** | **-0.092** | **0.006** | **-13.61** | **<0.001** |
| Num. Neighbours | **0.014** | **0.003** | **3.67** | **<0.001** |
| *Zero inflation model* | | | | |
| **Min Temp.** | **-1.174** | **0.431** | **-2.723** | **0.006** |
| **Min Temp.^2** | **1.195** | **0.426** | **2.806** | **0.005** |
| Laying date | 0.090 | 0.049 | 1.814 | 0.069 |
| Num. Neighbours | -0.012 | 0.032 | -0.369 | 0.712 |

SM-1 Table 3: Results from the hurdle models analysing the association between number of fledglings and temperature at the hatching period. The *conditional part* models the non-zero counts of our data and the *zero-inflated part* that models the probability of an observation being zero (i.e. the probability of brood failure). Bold denotes statistical significance. The model included mother identity and year as random factors.

| Variable | Estimate | Std. Error | z | P |
|---|---|---|---|---|
| *Temperatures at the hatching period (n=10421)* | | | | |
| Conditional model | | | | |
| **Mean Temp.** | **0.156** | **0.044** | **3.53** | **<0.001** |
| **Mean Temp.^$^2$** | **-0.141** | **0.043** | **-3.23** | **0.001** |
| **Laying date** | **-0.082** | **0.006** | **-12.79** | **<0.001** |
| **Num. Neighbours** | **0.014** | **0.003** | **3.69** | **<0.001** |
| *Zero inflation model* | | | | |
| **Mean Temp.** | **-1.114** | **0.349** | **-3.185** | **0.001** |
| **Mean Temp.^$^2$** | **1.055** | **0.342** | **3.081** | **0.002** |
| **Laying date** | **0.136** | **0.051** | **2.681** | **0.007** |
| Num. Neighbours | -0.010 | 0.032 | -0.324 | 0.745 |

SM-1 Table 4: Results from the hurdle models analysing the association between number of fledglings and temperature at the nestling period. The *conditional part* models the non-zero counts of our data and the *zero-inflated part* that models the probability of an observation being zero (i.e. the probability of brood failure). Bold denotes statistical significance. The model included mother identity and year as random factors.

| Variable | Estimate | Std. Error | z | P |
|---|---|---|---|---|
| *Temperatures at the nestling period (n=10421)* | | | | |
| Conditional model | | | | |
| Mean Temp. | 0.050 | 0.045 | 1.10 | 0.270 |
| Mean Temp.^$^2$ | -0.027 | 0.046 | -0.61 | 0.544 |
| **Laying date** | **-0.087** | **0.006** | **-12.92** | **<0.001** |
| **Num. Neighbours** | **0.014** | **0.003** | **3.61** | **<0.001** |
| *Zero inflation model* | | | | |
| Min Temp. | -0.434 | 0.394 | -1.101 | 0.270 |
| Min Temp.^$^2$ | 0.285 | 0.339 | 0.714 | 0.475 |
| **Laying date** | **0.178** | **0.052** | **3.372** | **<0.001** |
| Num. Neighbours | -0.009 | 0.032 | -0.298 | 0.765 |

SM-1 Table 5: Results from the hurdle models analysing the association between number of fledglings and temperature at the fledging period. The *conditional part* models the non-zero counts of our data and the *zero-inflated part* that models the probability of an observation being zero (i.e. the probability of brood failure). Bold denotes statistical significance. The model included mother identity and year as random factors.

| Variable | Estimate | Std. Error | z | P |
|---|---|---|---|---|
| *Temperatures at the fledging period (n=10421)* | | | | |
| Conditional model | | | | |
| **Mean Temp.** | **0.193** | **0.058** | **3.31** | **<0.001** |
| **Mean Temp.^$^2$** | **-0.175** | **0.058** | **-3.02** | **0.002** |
| **Laying date** | **-0.082** | **0.006** | **-12.97** | **<0.001** |
| **Num. Neighbours** | **0.014** | **0.003** | **3.66** | **<0.001** |
| *Zero inflation model* | | | | |
| Min Temp. | -0.484 | 0.482 | -1.00 | 0.315 |
| Min Temp^$^2$. | 0.539 | 0.480 | 1.124 | 0.261 |
| Laying date | 0.086 | 0.048 | 1.766 | 0.077 |
| Num. Neighbours | -0.012 | 0.032 | -0.367 | 0.713 |

### Supplementary Material 2

*Outputs from the models linking average temperature in the periods relative to breeding and fledging success.*

SM-2 Table 1: Results from the generalised linear mixed model analysing the association between fledgling success and temperature at egg-laying. Bold denotes statistical significance. The model included mother identity and year as random factors.

| Variable | Estimate | Std. Error | z | P |
|---|---|---|---|---|
| *Temperatures at egg-laying (n=11033)* | | | | |
| **Mean Temp.** | **1.792** | **0.133** | **13.473** | **<0.001** |
| **Mean Temp.^2** | **-1.905** | **0.131** | **-14.508** | **<0.001** |
| Laying date | -0.057 | 0.030 | -1.872 | 0.061 |
| **Clutch size** | **-0.063** | **0.019** | **-3.353** | **<0.001** |
| Num. Neighbours | 0.007 | 0.016 | 0.490 | 0.623 |

SM-2 Table 2: Results from the generalised linear mixed model analysing the association between fledgling success and temperature at incubation. Bold denotes statistical significance. The model included mother identity and year as random factors.

| Variable | Estimate | Std. Error | z | P |
|---|---|---|---|---|
| *Temperatures at incubation (n=10421)* | | | | |
| **Mean Temp.** | **0.527** | **0.210** | **2.513** | **0.012** |
| **Mean Temp.^2** | **-0.511** | **0.207** | **-2.466** | **0.013** |
| **Laying date** | **-0.165** | **0.027** | **-5.980** | **<0.001** |
| **Clutch size** | **-0.084** | **0.019** | **-4.416** | **<0.001** |
| Num. Neighbours | 0.005 | 0.016 | 0.314 | 0.753 |

SM-2 Table 3: Results from the generalised linear mixed model analysing the association between fledgling success and temperature at the hatching period. Bold denotes statistical significance. The model included mother identity and year as random factors.

| Variable | Estimate | Std. Error | z | P |
|---|---|---|---|---|
| *Temperatures at the hatching period (n=10421)* | | | | |
| **Mean Temp.** | **1.198** | **0.158** | **7.571** | **<0.001** |
| **Mean Temp^2.** | **-1.172** | **0.156** | **-7.477** | **<0.001** |
| **Laying date** | **-0.185** | **0.027** | **-6.800** | **<0.001** |
| **Clutch size** | **-0.091** | **0.019** | **-4.788** | **<0.001** |
| Num. Neighbours | 0.005 | 0.016 | 0.356 | 0.722 |

SM-2 Table 4: Results from the generalised linear mixed model analysing the association between fledgling success and temperature at the nestling period. Bold denotes statistical significance. The model included mother identity and year as random factors.

| Variable | Estimate | Std. Error | z | P |
|---|---|---|---|---|
| *Temperatures at the nestling period (n=10421)* | | | | |
| Mean Temp. | 0.052 | 0.168 | 0.310 | 0.757 |
| Mean Temp^2. | -0.014 | 0.171 | -0.087 | 0.931 |
| **Laying date** | **-0.178** | **0.027** | **-6.468** | **<0.001** |
| **Clutch size** | **-0.088** | **0.019** | **-4.629** | **<0.001** |
| Num. Neighbours | 0.005 | 0.016 | 0.326 | 0.745 |

SM-2 Table 5: Results from the generalised linear mixed model analysing the association between fledgling success and temperature at the fledging period. Bold denotes statistical significance. The model included mother identity and year as random factors.

| Variable | Estimate | Std. Error | z | P |
|---|---|---|---|---|
| *Temperatures at the fledging period (n=10421)* | | | | |
| **Mean Temp.** | **0.906** | **0.207** | **4.372** | **<0.001** |
| **Mean Temp^$^2$.** | **-0.961** | **0.207** | **-4.643** | **<0.001** |
| **Laying date** | **-0.156** | **0.026** | **-5.911** | **<0.001** |
| **Clutch size** | **-0.085** | **0.019** | **-4.491** | **<0.001** |
| Num. Neighbours | 0.007 | 0.016 | 0.429 | 0.668 |

**Supplementary Material 3**

*Outputs from the models linking average temperature in the periods relative to breeding mismatch with the winter-moth half-fall dates.*

To build Figure 3 in the main text considering the errors around our estimations of the temperature that maximises fledgling success, minimises the probability of brood failure and maximises number of fledglings we fitted models with the same structure than those presented above but following a Bayesian approach. For fledging success, we fitted generalised linear mixed models (GLMM) with a *binomial* error distribution. For the probability of brood failure, we coded number of fledglings as zero when a breeding attempt had zero fledglings and as one when it had one or more than one fledgling. Here, we fitted GLMMs with a *Bernoulli* error distribution. Finally, for number of fledglings we fitted GLMMs with Poisson distribution of errors, and number of fledglings (>0) as dependent variable. In these models, we included mean temperature of the period as linear and quadratic, laying date, clutch size (only for fledgling success) and number of neighbours included as fixed effects. Mother identity and year were included as random effects. Models were run for 10,000 iterations across four chains with a warm-up of 2000 iterations and a thinning of 10 iterations to ensure convergence and good effective sample sizes.

We used the posterior distributions of the models to compute the median temperature that maximises fledging success and minimises the probability of brood failure (*optimal temperature*) in addition to the 95% credible intervals. Optimal temperatures were computed as [$-b / (2*a)$], where $b$ represents the coefficient for the linear and $a$ the quadratic term of temperature. The optimal temperatures obtained from the frequentist and Bayesian approaches yield almost identical results except in those periods were the association between temperature and fledgling success probability of brood failure was not significant (i.e. 95% CI included zero; SM4 – Table 1).

SM-3 Table 1 Optimal temperature values from frequentist and Bayesian approaches.

| Period | Optimal temperature from frequentist approach | Median optimal temperature from Bayesian approach | Lower 95% CI | Upper 95% CI | Median temperature in the period | Standard deviation of temperature in the period |
|---|---|---|---|---|---|---|
| | | | *Fledgling success* | | | |
| Egg-laying | 9.39 | 9.39 | 9.14 | 9.63 | 9.94 | 2.01 |
| Incubation | 11.33 | 11.29 | 9.53 | 13.85 | 10.89 | 1.52 |
| Hatching period | 12.19 | 12.19 | 11.72 | 12.65 | 11.51 | 1.87 |
| Nestling period | 79.70 | 13.57 | -28.21 | 68.13 | 12.74 | 1.86 |
| Fledging period | 13.20 | 13.20 | 12.34 | 13.87 | 13.44 | 1.74 |
| | | | *Probability of brood failure* | | | |
| Egg-laying | 9.23 | 9.23 | 8.13 | 10.14 | 9.94 | 2.01 |
| Incubation | 10.69 | 10.79 | 8.97 | 12.19 | 10.89 | 1.52 |
| Hatching period | 12.58 | 12.57 | 11.39 | 14.12 | 11.51 | 1.87 |
| Nestling period | 20.20 | 16.87 | -44.57 | 93.85 | 12.74 | 1.86 |
| Fledging period | 12.62 | 12.97 | -4.92 | 29.59 | 13.44 | 1.74 |
| | | | Number of fledglings | | | |
| Egg-laying | 10.60 | 10.61 | 10.27 | 10.96 | 9.93 | 1.98 |
| Incubation | 5.20 | 6.10 | -31.35 | 59.99 | 10.90 | 1.50 |
| Hatching period | 13.23 | 13.20 | 11.99 | 15.10 | 11.53 | 1.87 |
| Nestling period | 24.18 | 17.87 | -72.97 | 120.9 | 12.73 | 1.86 |
| Fledging period | 15.42 | 15.41 | 14.15 | 17.57 | 13.41 | 1.74 |

**Supplementary Material 4**

*Outputs from the models linking average temperature in the periods relative to breeding mismatch with the winter-moth half-fall dates.*

SM-4 Table 1: Results from the linear mixed models analysing the association between absolute mismatch with winter-moth half-fall dates and temperature at the periods relative to breeding. Bold denotes statistical significance. The model included mother identity and year as random factors. Sample size is smaller in this set of models as half-fall dates were available for 45 of the 59 years.

| Variable | Estimate | Std. Error | z | P |
|---|---|---|---|---|
| *Temperatures at egg-laying (n=9711)* | | | | |
| **Mean Temp.** | **-4.552** | **0.170** | $\mathbf{F_{1,9479.0}=684.22}$ | **<0.001** |
| **Mean Temp^2.** | **0.217** | **0.008** | $\mathbf{F_{1,9486.1}=657.62}$ | **<0.001** |
| *Temperatures at incubation (n=9711)* | | | | |
| **Mean Temp.** | **-6.828** | **0.354** | $\mathbf{F_{1,7990.4}=372.04}$ | **<0.001** |
| **Mean Temp^2.** | **0.256** | **0.015** | $\mathbf{F_{1,8194.1}=258.30}$ | **<0.001** |
| *Temperatures at the hatching period (n=9711)* | | | | |
| **Mean Temp.** | **-4.543** | **0.022** | $\mathbf{F_{1,9414.9}=403.72}$ | **<0.001** |
| **Mean Temp^2.** | **0.165** | **0.009** | $\mathbf{F_{1,9437.0}=302.10}$ | **<0.001** |
| *Temperatures at the nestling period (n=9711)* | | | | |
| **Mean Temp.** | **-7.707** | **0.236** | $\mathbf{F_{1,9662.0}=1058.79}$ | **<0.001** |
| **Mean Temp^2.** | **0.266** | **0.009** | $\mathbf{F_{1,9643.4}=845.08}$ | **<0.001** |

**Supplementary Material 5**

*Outputs from the great tit models using the reduced dataset*

To explore the potential biases associated with using the reduced dataset (45 years instead of 59, see methods) for which winter moth data was available we repeated all the great tit analyses using the same modelling approaches and model structure but using this reduced dataset. The results from these analyses using the reduced dataset are qualitatively equal to those obtained from the full data.

1. *Temporal trends in temperature using the fixed time interval*

SM-5 Table 1: Result from the linear model analysing the temporal trends in mean temperature during the fixed interval with the reduced dataset.

| Variable | Estimate | Std. Error | z | P |
|---|---|---|---|---|
| | | *Fixed window* | | |
| Mean Temp. | 0.035 | 0.006 | $F_{1,43}$=25.21 | <0.001 |

2. *Temporal trends in temperature using the relative time intervals*

SM-5 Table 2: Result from the linear mixed models analysing the temporal trends in temperature in the five periods defined relative to individual reproductive timing using the reduced dataset. Egg-laying represents the period between laying the first and last eggs and incubation the period from the end of egg-laying to hatching. The periods hatching, nestling and fledging capture the time between hatching and 24 days post-hatch in 8-day intervals. The models included female individual identity and year (as a categorical variable) as random effects.

| **Reproductive Period** | **Estimate** | **SE** | ***F*** | ***P*** |
|---|---|---|---|---|
| | | *Egg-laying* | | |
| ***Year*** | -0.012 | 0.010 | $F_{1,43.2}$=1.308 | 0.258 |
| | | *Incubation* | | |
| ***Year*** | -0.007 | 0.011 | $F_{1,43.0}$=0.400 | 0.529 |
| | | *Hatching period* | | |
| ***Year*** | 0.0004 | 0.010 | $F_{1,43}$=0.002 | 0.964 |
| | | *Nestling period* | | |
| ***Year*** | -0.006 | 0.010 | $F_{1,43.1}$=0.380 | 0.540 |
| | | *Fledging period* | | |
| ***Year*** | -0.012 | 0.011 | $F_{1,43.0}$=1.669 | 0.286 |

#### 3. *Fitness consequences of mean temperatures in the relative periods*

##### *Number of fledglings*

SM-5 Table 3: Results from the hurdle models analysing the association between number of fledglings and temperature at egg-laying using the reduced dataset. The *conditional part* models the non-zero counts of our data and the *zero-inflated part* that models the probability of an observation being zero (i.e. the probability of brood failure). Bold denotes statistical significance. The model included mother identity and year as random factors.

| Variable | Estimate | Std. Error | z | P |
|---|---|---|---|---|
| *Temperatures at egg-laying* | | | | |
| Conditional model | | | | |
| **Mean Temp.** | **0.424** | **0.046** | **9.20** | **<0.001** |
| **Mean Temp^2.** | **-0.403** | **0.046** | **-8.74** | **<0.001** |
| **Laying date** | **-0.099** | **0.008** | **-12.23** | **<0.001** |
| **Num. Neighbours** | **0.018** | **0.004** | **4.22** | **<0.001** |
| *Zero inflation model* | | | | |
| **Min Temp.** | **-1.019** | **0.279** | **-3.644** | **<0.001** |
| **Min Temp^2.** | **1.087** | **0.271** | **3.997** | **<0.001** |
| Laying date | 0.079 | 0.054 | 1.469 | 0.141 |
| Num. Neighbours | -0.008 | 0.030 | -0.268 | 0.788 |

SM-5 Table 4: Results from the hurdle models analysing the association between number of fledglings and temperature at incubation using the reduced dataset. The *conditional part* models the non-zero counts of our data and the *zero-inflated part* that models the probability of an observation being zero (i.e. the probability of brood failure). Bold denotes statistical significance. The model included mother identity and year as random factors.

| Variable | Estimate | Std. Error | z | P |
|---|---|---|---|---|
| *Temperatures at incubation* | | | | |
| Conditional model | | | | |
| Mean Temp. | -0.091 | 0.070 | -1.30 | 0.193 |
| Mean Temp.^2 | 0.130 | 0.069 | 1.87 | 0.061 |
| **Laying date** | **-0.093** | **0.007** | **-12.70** | **<0.001** |
| Num. Neighbours | **0.018** | **0.004** | **4.41** | **<0.001** |
| *Zero inflation model* | | | | |
| Min Temp. | -0.905 | 0.465 | -1.945 | 0.051 |
| **Min Temp.^2** | **0.935** | **0.462** | **2.024** | **0.042** |
| Laying date | 0.097 | 0.050 | 1.922 | 0.054 |
| Num. Neighbours | -0.005 | 0.033 | -0.173 | 0.862 |

SM-5 Table 5: Results from the hurdle models analysing the association between number of fledglings and temperature at the hatching period using the reduced dataset. The *conditional part* models the non-zero counts of our data and the *zero-inflated part* that models the probability of an observation being zero (i.e. the probability of brood failure). Bold denotes statistical significance. The model included mother identity and year as random factors.

| Variable | Estimate | Std. Error | z | P |
|---|---|---|---|---|
| *Temperatures at the hatching period* | | | | |
| Conditional model | | | | |
| **Mean Temp.** | **0.206** | **0.051** | **4.04** | **<0.001** |
| **Mean Temp.^2** | **-0.193** | **0.050** | **-3.85** | **<0.001** |
| **Laying date** | **-0.090** | **0.007** | **-12.25** | **<0.001** |
| **Num. Neighbours** | **0.019** | **0.004** | **4.49** | **<0.001** |
| *Zero inflation model* | | | | |
| **Mean Temp.** | **-1.098** | **0.360** | **-3.047** | **0.002** |
| **Mean Temp.^2** | **1.036** | **0.346** | **2.988** | **0.002** |
| **Laying date** | **0.140** | **0.053** | **2.638** | **0.008** |
| Num. Neighbours | -0.003 | 0.033 | -0.115 | 0.908 |

SM-5 Table 6: Results from the hurdle models analysing the association between number of fledglings and temperature at the nestling period using the reduced dataset. The *conditional part* models the non-zero counts of our data and the *zero-inflated part* that models the probability of an observation being zero (i.e. the probability of brood failure). Bold denotes statistical significance. The model included mother identity and year as random factors.

| Variable | Estimate | Std. Error | z | P |
|---|---|---|---|---|
| *Temperatures at the nestling period* | | | | |
| Conditional model | | | | |
| Mean Temp. | 0.117 | 0.051 | 2.29 | 0.022 |
| Mean Temp.^2 | -0.091 | 0.051 | -1.76 | 0.079 |
| **Laying date** | **-0.102** | **0.008** | **-12.34** | **<0.001** |
| **Num. Neighbours** | **0.018** | **0.004** | **4.40** | **<0.001** |
| *Zero inflation model* | | | | |
| Min Temp. | -0.612 | 0.397 | -1.542 | 0.123 |
| Min Temp.^2 | 0.491 | 0.395 | 1.244 | 0.213 |
| **Laying date** | **0.182** | **0.057** | **3.166** | **0.001** |
| Num. Neighbours | -0.003 | 0.033 | -0.101 | 0.919 |

SM-5 Table 6: Results from the hurdle models analysing the association between number of fledglings and temperature at the fledgling period using the reduced dataset. The *conditional part* models the non-zero counts of our data and the *zero-inflated part* that models the probability of an observation being zero (i.e. the probability of brood failure). Bold denotes statistical significance. The model included mother identity and year as random factors.

| Variable | Estimate | Std. Error | z | P |
|---|---|---|---|---|
| *Temperatures at the fledging period* | | | | |
| Conditional model | | | | |
| **Mean Temp.** | **0.162** | **0.069** | **2.34** | **0.019** |
| **Mean Temp.^2** | **-0.141** | **0.069** | **-2.03** | **0.042** |
| **Laying date** | **-0.090** | **0.007** | **-12.42** | **<0.001** |
| **Num. Neighbours** | **0.018** | **0.004** | **4.40** | **<0.001** |
| *Zero inflation model* | | | | |
| Min Temp. | -7.09 | 0.552 | -1.358 | 0.174 |
| Min Temp^2. | 0.748 | 0.520 | 1.438 | 0.150 |
| Laying date | 0.097 | 0.050 | 1.923 | 0.054 |
| Num. Neighbours | -0.005 | 0.033 | -0.148 | 0.882 |

*Fledgling success*

SM-5 Table 7: Results from the generalised linear mixed model analysing the association between fledgling success and temperature at egg-laying using the reduced dataset. Bold denotes statistical significance. The model included mother identity and year as random factors.

| Variable | Estimate | Std. Error | z | P |
|---|---|---|---|---|
| *Temperatures at egg-laying* | | | | |
| **Mean Temp.** | **2.059** | **0.149** | **13.75** | **<0.001** |
| **Mean Temp.^2** | **-2.164** | **0.148** | **-14.601** | **<0.001** |
| Laying date | **-0.106** | **0.033** | **-3.130** | **0.001** |
| **Clutch size** | **-0.129** | **0.021** | **-6.135** | **<0.001** |
| Num. Neighbours | 0.017 | 0.017 | 0.987 | 0.323 |

SM-5 Table 8: Results from the generalised linear mixed model analysing the association between fledgling success and temperature at incubation using the reduced dataset. Bold denotes statistical significance. The model included mother identity and year as random factors.

| Variable | Estimate | Std. Error | z | P |
|---|---|---|---|---|
| *Temperatures at incubation* | | | | |
| Mean Temp. | 0.128 | 0.237 | 0.540 | 0.589 |
| Mean Temp.^2 | -0.098 | 0.235 | -0.418 | 0.676 |
| **Laying date** | **-0.195** | **0.030** | **-6.448** | **<0.001** |
| **Clutch size** | **-0.125** | **0.021** | **-5.955** | **<0.001** |
| Num. Neighbours | 0.010 | 0.018 | 0.578 | -0.563 |

SM-5 Table 9: Results from the generalised linear mixed model analysing the association between fledgling success and temperature at the hatching period using the reduced dataset. Bold denotes statistical significance. The model included mother identity and year as random factors.

| Variable | Estimate | Std. Error | z | P |
|---|---|---|---|---|
| *Temperatures at the hatching period* | | | | |
| **Mean Temp.** | **1.048** | **0.173** | **6.037** | **<0.001** |
| **Mean Temp^2.** | **-0.997** | **0.170** | **-5.866** | **<0.001** |
| **Laying date** | **-0.218** | **0.030** | **-7.120** | **<0.001** |
| **Clutch size** | **-0.131** | **0.021** | **-6.220** | **<0.001** |
| Num. Neighbours | 0.010 | 0.018 | 0.563 | 0.574 |

SM-5 Table 10: Results from the generalised linear mixed model analysing the association between fledgling success and temperature at the nestling period using the reduced dataset. Bold denotes statistical significance. The model included mother identity and year as random factors.

| Variable | Estimate | Std. Error | z | P |
|---|---|---|---|---|
| *Temperatures at the nestling period* | | | | |
| **Mean Temp.** | **0.393** | **0.185** | **2.127** | **0.033** |
| Mean Temp^2. | -0.359 | 0.185 | -1.936 | 0.052 |
| **Laying date** | **-0.213** | **0.032** | **-6.500** | **<0.001** |
| **Clutch size** | **-0.127** | **0.021** | **-6.034** | **<0.001** |
| Num. Neighbours | 0.010 | 0.018 | 0.589 | 0.555 |

SM-2 Table 11: Results from the generalised linear mixed model analysing the association between fledgling success and temperature at the fledging period using the reduced dataset. Bold denotes statistical significance. The model included mother identity and year as random factors.

| Variable | Estimate | Std. Error | z | P |
|---|---|---|---|---|
| *Temperatures at the fledging period* | | | | |
| **Mean Temp.** | **1.381** | **0.242** | **5.709** | **<0.001** |
| **Mean Temp^$^{2}$.** | **-1.414** | **0.242** | **-5.833** | **<0.001** |
| **Laying date** | **-0.190** | **0.029** | **-6.400** | **<0.001** |
| **Clutch size** | **-0.125** | **0.021** | **-5.977** | **<0.001** |
| Num. Neighbours | 0.011 | 0.018 | 0.634 | 0.526 |

**Supplementary Material 6**
*Sliding window analysis*

We used the R package *climwin* (v. 1.2.3; Bailey, L.D. (2016) to estimate the window where mean temperature was most strongly associated with the onset of reproduction in our population. We used the same model specifications as in Simmonds et al. (2019) who successfully used this approach in a subset of our data. We used absolute sliding time windows, with the 20$^{th}$ of May as reference day. We allowed windows to vary 365 days in length. We found that the temperature window that best explained the onset of reproduction in our data started the 15$^{th}$ of February and ended the 2$^{nd}$ of May. We used the critical probability to check whether the window we detected was just by chance. With that aim, we ran 5 randomised models using different randomisations of the onset of reproduction (van de Pol, M. 2016). We used these randomisations and the original model to compute the critical probability which was $p<0.001$. Smaller p-values are associated with a smaller probability of the window we detected was by chance.

### **Supplementary Material 7**

*Computing mean temperatures relative to each breeding attempt*

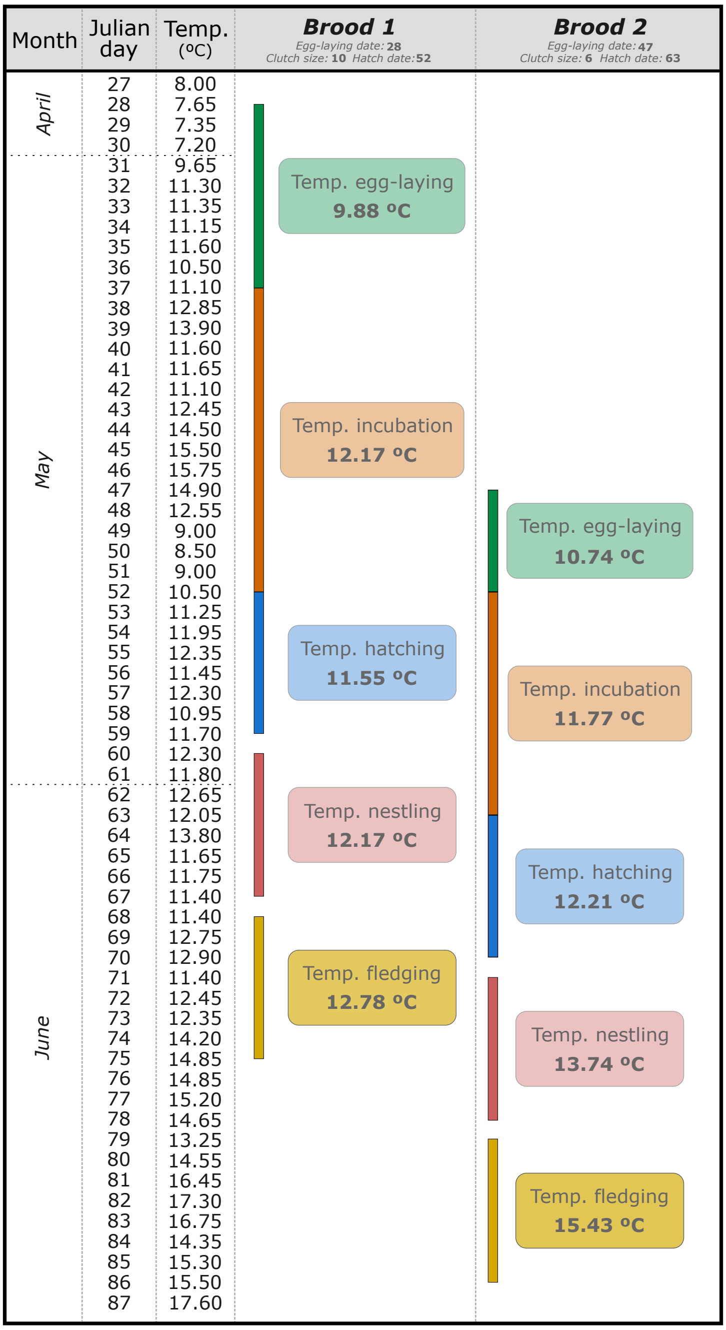


SM-7 Figure 1: Illustrative figure of the used followed to compute average temperature during periods relative to the timing of each individual breeding attempt, using two broods of the year 1988 as an example. Julian date represents the number of days since April 1$^{st}$, and Temp. ($^{o}$ C) represents the average temperature each day. Green represents the egg-laying period, the interval between laying the first and last egg. Orange represents the incubation period, the interval between laying the last egg and hatching. Blue represents the hatching period, the interval between hatching and 7-days post-hatch. Red represents the nestling period, the interval between 8- and 15-days post-hatch. Grey represents the fledging period, the interval between 16- and 23-days post-hatch.

**Supplementary Material 8**

*Temporal trends in temperature around half-fall with different window sizes*

SM-8 Table 1: Results from the linear models exploring the temporal trends of temperature around half-fall using different size windows.

| Variable | Estimate | Std. Error | z | P |
|---|---|---|---|---|
| | | *±7 days around half-fall* | | |
| Mean Temp. | -0.003 | 0.011 | $F_{1,43}$=0.119 | 0.731 |
| | | *±15 days around half-fall* | | |
| Mean Temp. | -0.002 | 0.008 | $F_{1,43}$=0.060 | 0.807 |
| | | *±20 days around half-fall* | | |
| Mean Temp. | -0.003 | 0.006 | $F_{1,43}$=0.264 | 0.610 |

**Supplementary Material 9**

*Temperature during the relative periods and raw mismatch with peak of caterpillar availability*

To analyse the association between mismatch and temperature in the relative windows we repeated the linear mixed models presented in the main text but using the raw values of mismatch as dependent variable instead of the absolute values. As dependent variables we included the temperature of the period and its quadratic component as explanatory. In these models we included year (as a categorical variable) and female identity as random effects.

Our results show significant negative associations between mismatch and temperature in the four periods (Table SM-9 Table 1). With above and below optimum temperatures increasing leading to broods being late or early respectively to the perfect match between the peak of food availability and demands.

SM-9 Table 1: Results from the linear mixed models analysing the association between absolute mismatch with winter-moth half-fall dates and temperature at the periods relative to breeding. Bold denotes statistical significance. The model included mother identity and year as random factors. Sample size is smaller in this set of models as half-fall dates were available for 45 of the 59 years.

| Variable | Estimate | Std. Error | z | P |
|---|---|---|---|---|
| *Temperatures at egg-laying (n=9711)* | | | | |
| **Mean Temp.** | **-1.331** | **0.027** | **$F_{1,9176.5}$=2397.3** | **<0.001** |
| *Temperatures at incubation (n=9711)* | | | | |
| **Mean Temp.** | **-2.027** | **0.056** | **$F_{1,8178.9}$=1271.2** | **<0.001** |
| *Temperatures at the hatching period (n=9711)* | | | | |
| **Mean Temp.** | **-1.428** | **0.033** | **$F_{1,9142.9}$=1860.4** | **<0.001** |
| *Temperatures at the nestling period (n=9711)* | | | | |
| **Mean Temp.** | **-1.949** | **0.028** | **$F_{1,9613}$=4674.0** | **<0.001** |

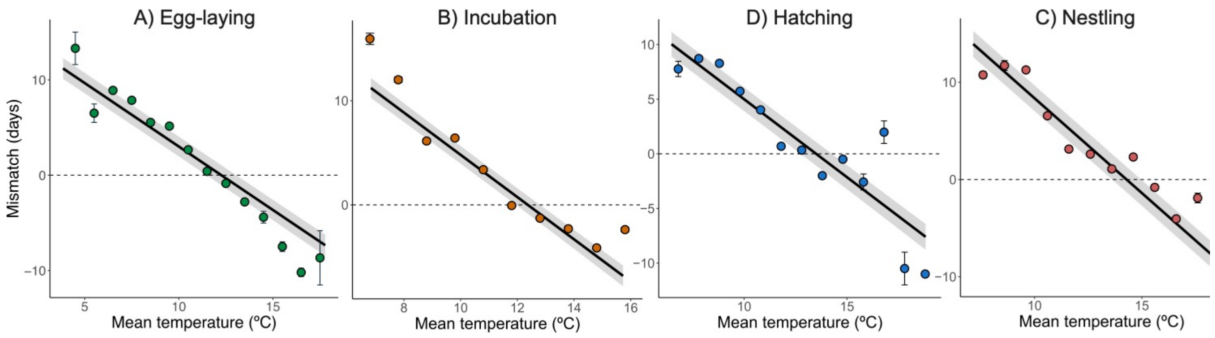


SM 9 – Figure 1. Associations between raw mismatch with the peak of food availability (half-fall date) and mean temperature (º C) in the windows relative to individual breeding. The solid line and grey ribbon represent the predicted association and 95% confidence interval from the model. The coloured dots and lines represent the average brood failure probability ± standard error grouped in 1° C bins. The dotted horizontal line (y=0) represents a perfect match between the peak of food availability and demand. The fledging period was not included as it occurs after the half-fall date. The numbers above dots denote the number of observations.
